# Screening for highly transduced genes in *Staphylococcus aureus* reveals both lateral and specialized transduction

**DOI:** 10.1101/2020.10.28.360172

**Authors:** Janine Zara Bowring, Yue Su, Ahlam Alsaadi, Sine L. Svenningsen, Julian Parkhill, Hanne Ingmer

## Abstract

Bacteriophage-mediated transduction of bacterial DNA is a major route of horizontal gene transfer in the human pathogen, *Staphylococcus aureus*. Transduction involves packaging of bacterial DNA by viruses and enables transmission of virulence and resistance genes between cells. To learn more about transduction in *S. aureus*, we searched a transposon mutant library for genes and mutations that enhanced transfer mediated by the temperate phage, φ11. Using a novel screening strategy, we performed multiple rounds of transduction of transposon mutant pools selecting for an antibiotic resistance marker within the transposon element. When determining the locations of transferred mutations, we found that, within each pool of 96 mutants the screen had selected for just 1 or 2 transposon mutant(s). Subsequent analysis showed that the position of the transposon, rather than inactivation of bacterial genes, was responsible for the phenotype. Interestingly, from multiple rounds we identified a pattern of transduction that encompassed mobile genetic elements, as well as chromosomal regions both upstream and downstream of the phage integration site. The latter was confirmed by DNA sequencing of purified phage lysates. Importantly, transduction frequencies were lower for phage lysates obtained by phage infection rather than induction. Our results confirm previous reports of lateral transduction of bacterial DNA downstream of the integrated phage, but also indicate specialized transduction of DNA upstream of the phage, likely involving imprecise excision of the phage from the bacterial genome. These findings illustrate the complexity of transduction processes and increase our understanding of the mechanisms by which phages transfer bacterial DNA.

**Importance:** Horizontal transfer of DNA between bacterial cells contributes to the spread of virulence and antibiotic resistance genes in human pathogens. For *Staphylococcus aureus*, bacterial viruses are particularly important. These viruses, termed bacteriophages, can transfer bacterial DNA between cells by a process known as transduction, which despite of its importance is only poorly characterized. Here, we employed a transposon mutant library to investigate transduction in *S. aureus*. We show that the location of bacterial DNA in relation to bacteriophages integrated in the bacterial genome is a key decider of how frequently that DNA is transduced. Based on serial transduction of transposon mutant pools and direct sequencing of bacterial DNA in bacteriophage particles, we demonstrate both lateral and specialized transduction. The use of mutant libraries to investigate the patterns of bacterial DNA transfer between cells could help understand how bacteria evolve virulence and resistance and may ultimately lead to new intervention strategies.

## Introduction

*S. aureus* is a gram-positive opportunistic pathogen that causes a wide-range of disease in humans. It can become resistant to a variety of antibiotics like methicillin (MRSA), tetracycline and vancomycin, and resistance is limiting treatment options (1, 2). The staphylococcal genome is highly plastic, largely due to horizontal gene transfer mediated by mobile genetic elements (MGEs) such as staphylococcal pathogenicity islands (SaPIs), plasmids and phages (3, 4). Particularly, phage mediated transduction appears to be an important route of gene transfer. Transduction is a process whereby some phages are able to package bacterial DNA at low frequency and transfer this DNA between cells (5, 6). Transducing phages are often temperate as they can both be integrated in the bacterial genome as prophages (lysogeny) or be lytic, where they replicate before lysing the cell and releasing the progeny phages. Transduction can occur through a number of different mechanisms; generalised transduction, specialised transduction, and lateral transduction. In *S. aureus*, generalized transduction has been used for many years as a genetic tool and recently, the mechanism for lateral transduction was established (7, 8). In contrast we know very little about specialized transduction in this organism.

The often-used *S. aureus* laboratory strain 8325-4 is the phage-cured equivalent of strain 8325, which contains three prophages (φ11, φ12, and φ13) ((9). Phage φ11 is one of the best characterised of the staphylococcal phages, and regularly used as a laboratory tool for transferring genes by transduction (10, 11). As a prophage, it sits in an intergenic region of the 8325 chromosome with a specific directionality, where the att_L_ is situated at ~1.967 Mb and the att_R_ at ~1.923 Mb (12, 13). Phage φ11 is a pac type phage, meaning that it recognises a phage *‘pac’* site and packages DNA into the capsid in a ‘headful’ manner, terminating when the capsid is filled with slightly more than the ~45 kb phage genome (14). As such, φ11 can transfer bacterial DNA by generalised transduction, which occurs when bacterial DNA contains sequences that have homology to the phage *‘pac’* sites and so are recognised by the phage packaging machinery (14). Generalised transduction can occur either when a phage infects a bacterium and enters the lytic cycle, or when a resident prophage is induced.

Lateral transduction is the most recently identified form of phage-mediated transduction in *S. aureus* (14). Previously, it was thought that prophage induction initiated the excision, replication, and packaging cycle in that order, with replication of the phage only occurring post-excision. However, it has since been determined that in *S. aureus* some phages begin replication before excising from the host chromosome (14). This leads to the replication of the integrated prophage together with the flanking regions of bacterial DNA. Because the flanking bacterial DNA is replicated with the integrated prophage, the bacterial DNA downstream of the *pac* site will be packaged by the headful mechanism. In practice, this has led to transduction frequencies of 1000-fold higher rates when compared to generalised transduction rates and to higher rates of transduction up to 100 kb downstream of the prophage. The study identified that this effect was unidirectional from the pac site and so the region upstream of the phage was not transferred (15). Lateral transduction has only been shown to occur with *pac* prophages that have been induced and does not occur when a phage infects and undergoes the lytic cycle (16, 17).

Specialised transduction occurs when a prophage excises incorrectly from the bacterial chromosome (16–24). Instead of a precise excision of the full phage genome between the att*L* and att*R* sites formed at integration, an incomplete portion of the phage excises alongside some of the flanking bacterial DNA both upstream and downstream of the phage. As long as the phage *pac* site is present, this hybrid DNA will be replicated, packaged into phage capsids, and transduced (15, 17, 20). Specialised transduction only occurs when a prophage is induced, rather than when a phage infects and undergoes the lytic cycle. It is limited to the immediate regions of bacterial DNA flanking a prophage. Specialised transduction mechanisms have been reported for phages infecting a number of bacterial genera, including *Bacillus, Salmonella, Pseudomonas* and *Vibrio* (14, 25). However, there is an absence of literature on specialised transduction by staphylococcal phages, with most φ11 papers referring to the work on *Salmonella* phage P22 when describing the process of specialised transduction (25).

Different bacteria and the prophages display different patterns of transduction (26). This means that different strains within the same species, but that contain differing prophages, can have varying patterns of transduction. A recent bioRxiv preprint described the use of DNA short- and long-read sequencing to investigate the patterns of bacterial DNA in transductant particles between different genera and species of bacteria (27). This pattern could be explained by different phages and prophages being more or less prone to lateral, specialised or generalised transduction and by the number of pseudo-pac sites in the strain’s chromosome. Furthermore, the same temperate phage may transduce bacterial DNA differently depending on whether that phage is undergoing lytic infection or lysogenic induction. Both lateral and specialized transduction require the induction of an integrated prophage, as both are initially reliant on the phage being integrated into the bacterial chromosome. Generalised transduction, on the other hand, can occur during both life cycles.

To further understand transduction in *S. aureus*, we explored a screening methodology that allowed us to identify *S. aureus* transposon mutants that were preferentially transduced from a pool of mutants. We utilised the Nebraska transposon mutant library (NTML) in *S. aureus* strain JE2, infecting pools of 96 mutants with the transducing, lysogenic phage φ11 (27). The resultant lysates were transduced to recipient strain 8325-4 φ11, from which pooled colonies could be induced by mitomycin C and transduced in cycles, selecting for transductants with the transposon-encoded resistance marker. From this screen we anticipated identifying genes that either, when inactivated, promoted transduction, or were located within the chromosome in regions transduced at higher rates than elsewhere. Our results revealed location-specific effects of the highly transduced regions flanking the phage φ11 integration site, by comparing the transduction frequency of other genes located in the same region. Indeed, the regions flanking the φ11 integration site were further confirmed as preferentially transduced regions of bacterial DNA by sequencing purified phage lysates.

Here, we show that the transposon mutants in the chromosomal regions were transduced by both lateral and specialised transduction. This study establishes a simple screening method that reveals patterns of transduction at the genome-level, which can be readily applied to transposon mutant libraries of other bacteria. Remarkably, this study is the first to demonstrate specialised transduction in *S. aureus*.

## Materials and Methods

### Bacterial strains and growth conditions

Bacterial strains used in this study are listed in supplementary table 1. *S. aureus* strains were grown in TSB and on TSA plates (Oxoid), with or without antibiotics as appropriate.

### Bacteriophage titre

Phage titre quantifications were performed essentially as previously described (27). Briefly, recipient strains were grown to 0.35 OD600 and 100 μL aliquots of recipient mixed with 100 μL phage lysate, at different dilutions in phage buffer (10^0^ – 10^-8^; MgSO_4_ 1 mM, CaCl_2_ 4 mM, Tris-HCl pH 8 50 mM, NaCl_2_ 0.1 M). After 10 minutes incubation at room temperature, 3 mL of liquid PTA (phage top agar; Oxoid nutrient broth n°2, agar 3.5% w/v) was added and the mixture poured out on PB plates (phage base; nutrient broth n°2, agar 7% w/v). Plates were incubated at 37°C overnight and plaques counted.

### Bacteriophage transduction

Phage transductions were performed as previously described (27). Briefly, recipient strains were grown to 1.4 OD_600_, 4.4 mM CaCl_2_ added and 1 mL aliquots of recipient mixed with 100 μL phage lysate, at different dilutions in phage buffer (10^0^ – 10^-4^). After 20 minutes incubation at 37°C, 3 mL of liquid TTA (transduction top agar; TSA, 50% agar) was added and the mix poured out on selective TSA plates. Plates were incubated at 37°C for 24 hours and colonies counted.

### Bacteriophage infection and induction

Bacteriophage infection was performed as previously described (26, 28). Briefly, recipient bacteria were grown to 0.15 OD_600_ and centrifuged, before re-suspending the pellet in 1:1 TSB and phage buffer. The culture was then infected with phage at an MOI of 1 and incubated at 30°C, 80 rpm until complete visual lysis, normally after 3 – 4 hours. Lysates were then filtered using 0.22 μM filters and stored at 4°C.

Bacteriophage induction was performed as previously described (29). Briefly, lysogens were grown to 0.1 – 0.2 OD_600_ and 2 μg ml^-1^ mitomycin C (Sigma, from *Streptomyces caespitosus*) added to induce resident prophages. Cultures were then incubated at 30°C, 80 rpm until complete visual lysis, normally after 3 – 4 hours. Lysates were filtered using 0.22 μM filters and stored at 4°C.

### NTML transposon transduction screening

For screening plates from the NTML, one 96-well plate was replicated and grown overnight in selective TSB (erythromycin, 10 μg ml^-1^) before pooling 10 μL of each mutant in 100 mL selective TSB, in a 500 mL flask. The culture was infected with φ11 (MOI 1) as described. The resulting lysate was then used to transduce the NTML transposons to 8325-4 φ11 recipient, as described. The plate resulting from the transduction of non-diluted lysate was harvested, pooled and grown in selective TSB and the culture was diluted then induced. The resulting lysate was used to transduce into the 8325-4 φ11 recipient a second time, and the transductants were pooled and induced. From the resulting lysate, a third transduction was performed, 10 transductant colonies selected and the location of the transposon mutant identified as previously described (30).

### DNA sequencing

Preparation of the phage lysates for DNA sequencing was done by precipitating the phage capsids before DNA extraction. For the phage precipitation, DNase (2.5 U ml^-1^, Thermo) and RNase (1 μg ml^-1^, Sigma) were added to the filtered lysate before incubation at 37°C for 1 hour. NaCl_2_ (1M) was added and mixed on ice with an orbital shaker at 4°C for 1 hour. The lysate was then centrifuged at 11,000 x g for 10 minutes at 4°C and supernatant collected and mixed well with PEG 8000 (10% w/v) before incubation on ice overnight at 4°C. The lysate was then centrifuged at 11,000 x g for 10 minutes at 4°C and the supernatant discarded, with falcon tubes inverted and dried. The pellet appears as two vertical white lines and was resuspended in 8 mL phage buffer per 500 mL lysate, with resuspension for 1 hour at 4°C. DNA extraction was then performed using the MagAttract HMW DNA Kit (Qiagen) as per manufacturer’s instructions. DNA concentration and absorbance peaks were checked using a Thermo NanoDrop 2000 and a Qubit Flex Fluorometer. A draft genome for 8325-4 was generated by sequencing chromosomal DNA on an Illumina miSeq following the manufacturer’s instructions. The sequence was assembled, and the contigs ordered using the Sanger Institute pipelines (31). Phage lysate DNA was sequenced in the same way, and the resulting reads were mapped to this genome using SMALT (26, 28) and visualised using Artemis (26, 28).

### DNA methods

General DNA manipulations were performed using standard procedures. Primers used in this study can be found in supplementary table 2. PCRs were performed using DreamTaq Green master mix (2x; Thermo) as per manufacturer’s instructions. PCRs for identification of transposon locations were performed as previously described (7, 9). PCR products were sequenced at Eurofins MWG Operon.

For identification of the presence of prophages in strains, a multiplex PCR for the main integrases of the staphylococcal phages was performed, as previously described (32). Initial screening of colonies was performed using 1 μL of template from a single colony resuspended in 50 μL ddH_2_O, boiled for 10 mins and on ice for 10 mins. Results were confirmed using DNA template (10 ng), extracted using GenElute^™^ Bacterial Genomic DNA Kit as per manufacturer’s instructions (Sigma-Aldrich).

For the long amplifications necessary for amplifying regions of specialised transduction, the Takara LA polymerase PCR kit was used as per manufacturer’s instructions with the following alteration, a 30 s annealing step was added at 55°C. Templates used were either bacterial DNA extracted with the GenElute™ Bacterial Genomic DNA Kit or phage DNA extracted from phage lysates that had been precipitated as previously described and then extracted using the GenElute™ Bacterial Genomic DNA Kit with an extended proteinase K incubation of 1 hour (Sigma-Aldrich). 50 ng of template was used for each PCR. The resulting products were run out on a 0.7% agarose gel at 30 V overnight.

### Statistical analyses

Data analyses were performed in GraphPad Prism 8 software. Specific statistical tests are indicated in the figure legends, where appropriate.

## Results

### Transduction selection preferentially identifies transposon mutants in MGEs and regions flanking prophage φ11

With the aim of identifying genes or mutations that increase transduction frequency, we screened the *S. aureus* JE2 NTML for mutations that were preferentially transduced. To achieve this, we carried out consecutive rounds of transduction on pools of library mutants by combining 96-well plates of transposon mutants and infecting these with the transducing temperate phage φ11, to produce an ‘infection’ phage lysate containing transducing particles. We then performed transduction using these lysates, with the laboratory strain 8325-4 φ11 as recipient. Having the φ11 prophage in the recipient prevents phage superinfection and thus increases survival of transductants. Furthermore, previous studies have shown that lysogenic strains can undergo autotransduction, where a lysogen spontaneously releases phages that infect nearby susceptible strains, resulting in transductants that can return useful bacterial DNA to the original lysogenic population (27, 33). Transductants carrying the transposon element were selected for by their resistance to erythromycin, which is encoded within the transposon. These transductants were then pooled and the φ11 prophage was induced to produce a ‘round 1 induction’ lysate. The transduction and induction steps were repeated to produce ‘round 2’ and ‘round 3’ transduction lysates. At each stage the CFU ml^-1^ and PFU ml^-1^ were recorded and the transduction frequency was calculated, controlling for phage variation (CFU ml^-1^/ PFU ml^-1^; Fig 1). Importantly, the transduction frequencies showed a general increase over the three rounds of transduction screening for the eight 96-well plates examined, demonstrating that our selection procedure had succeeded (Fig 1).

**Figure 1.**
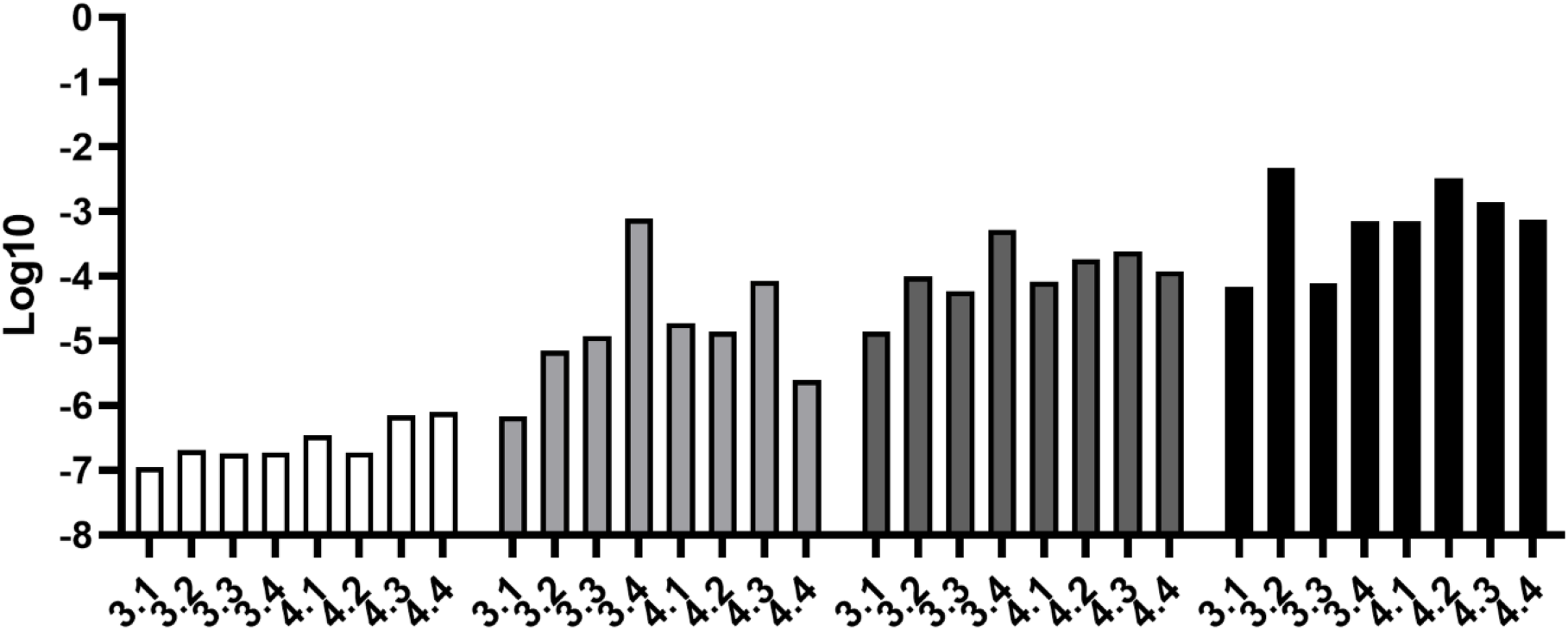
The transduction frequencies of pooled transposon mutants. Transduction frequencies (log transformed CFU ml^-1^/PFU ml^-1^) for the initial infection transduction (white), the rounds 1 (light grey), 2 (dark grey), and 3 (black) induction-transduction. CFU ml^-1^ was calculated from the number of transductant colonies on erythromycin selective plates and PFU ml^-1^ from the number of plaques formed on a susceptible recipient (8325-4). Each bar represents one replicate of a pool of 96 transposon mutants labelled by library plate number (3.1 – 4.4).

At the round 3 transduction, we collected 10 different colonies from each plate pool and identified the locations of the transposon mutations by digestion, ligation, and sequencing of the transposon-chromosomal flanks, using an established protocol (14). Examination of the 10 colonies from each of the different pools revealed that in the third round transductants, most pools contained just one predominant transposon mutant (Fig 2). For 7 of the 8 pools, only one transposon mutant was identified in the final round of colony check, whilst for plate 3-1, 2 transposon mutants were identified. The first striking observation was that the majority (5 out of 9) of the transposon mutants identified in our screen were situated on MGEs (Fig 2). Of these, one transposon was located in a SaPI5 gene, one in the USA300φSa2 prophage of JE2, while the other 3 were located in the USA300φSa3 JE2 prophage. These MGE-located transposon mutants were distributed throughout the chromosome (Fig 2). When we identified the locations of the other 4 chromosomally situated non-MGE related transposon mutants, we found that these clustered around the integration site of the φ11 (attB; Fig 2). Relative to the φ11 prophage directionality, 2 of the transposon mutants were situated upstream, but close to, the φ11 attB, whilst the other 2 were located downstream of this site (9, 14). We hypothesised that we were identifying transposon mutants that were preferentially transduced due to either induced MGE mobility or lateral and specialised transduction.

**Figure 2.**
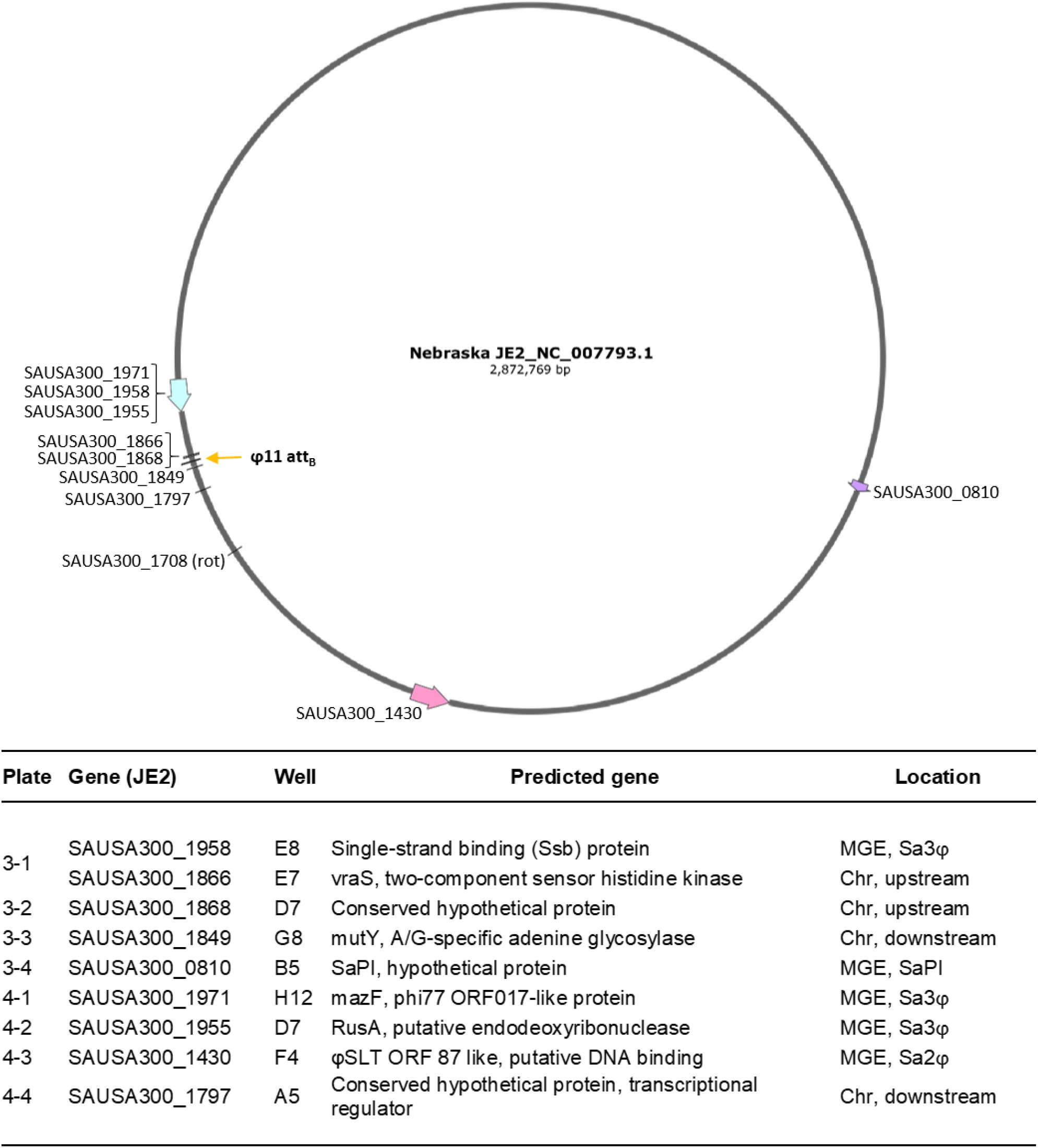
Locations of the identified transposon mutants in the JE2 genome. Depiction of NTML JE2 genome (NCBI accession number 007793.1), with MGEs USA300φSa2 (pink arrow), USA300φSa3 (blue arrow), and JE2 SaPI5 (purple arrow). The φ11 insertion site (att_B_) is indicated. The genes identified with a transposon mutant insertion are marked with the NTML gene identification number, as is the position of the Tn_rotB control. The table identifies the gene in which the transposon was inserted for each identified mutant. For all plates this was either 1 or 2 mutants. ‘Plate’ refers to the NTML 96-plate numbering of the pooled 96 mutants, ‘location’ identifies whether the transposon insertion was in a MGE-situated gene or in the chromosome (Chr). Upstream and downstream refer to the position of the gene relative to the φ11 prophage insertion, using the prophage directionality (eg. downstream = downstream of the att_R_, upstream = upstream of the att_L_).

### Transposon mutants up- and down-stream of φ11 have an increased transduction frequency

The chromosomal transposon mutants clustering around the φ11 att_B_ interested us as potential incidents of specialised or lateral transduction (SAUSA300_1866, SAUSA300_1868, SAUSA300_1849, and SAUSA300_1797). Therefore, we transduced these individual transposon mutants from JE2 into the clean laboratory strain 8325-4 φ11, which has only the φ11 prophage, and confirmed by “integrase PCR” that none of the JE2 endogenous phages had been transferred. Using these ‘clean background’ strains, we confirmed that the transduction frequencies in the 4 chromosomal transposon mutants were higher than our 8325-4 φ11 rotB::erm transposon control. The rotB transposon mutant has a wild-type phenotype in respect to bacterial growth and transduction, and φ11 should only transfer this transposon mutant by generalised transduction, based on its position in the 8325 genome (1.794 Mb, figure 2). All 4 transposon mutants had a higher transduction frequency than the rotB control, with mutants SAUSA300_1868, SAUSA300_1866, SAUSA300_1849, and SAUSA300_1797 having significantly higher frequencies (Fig 3).

**Figure 3.**
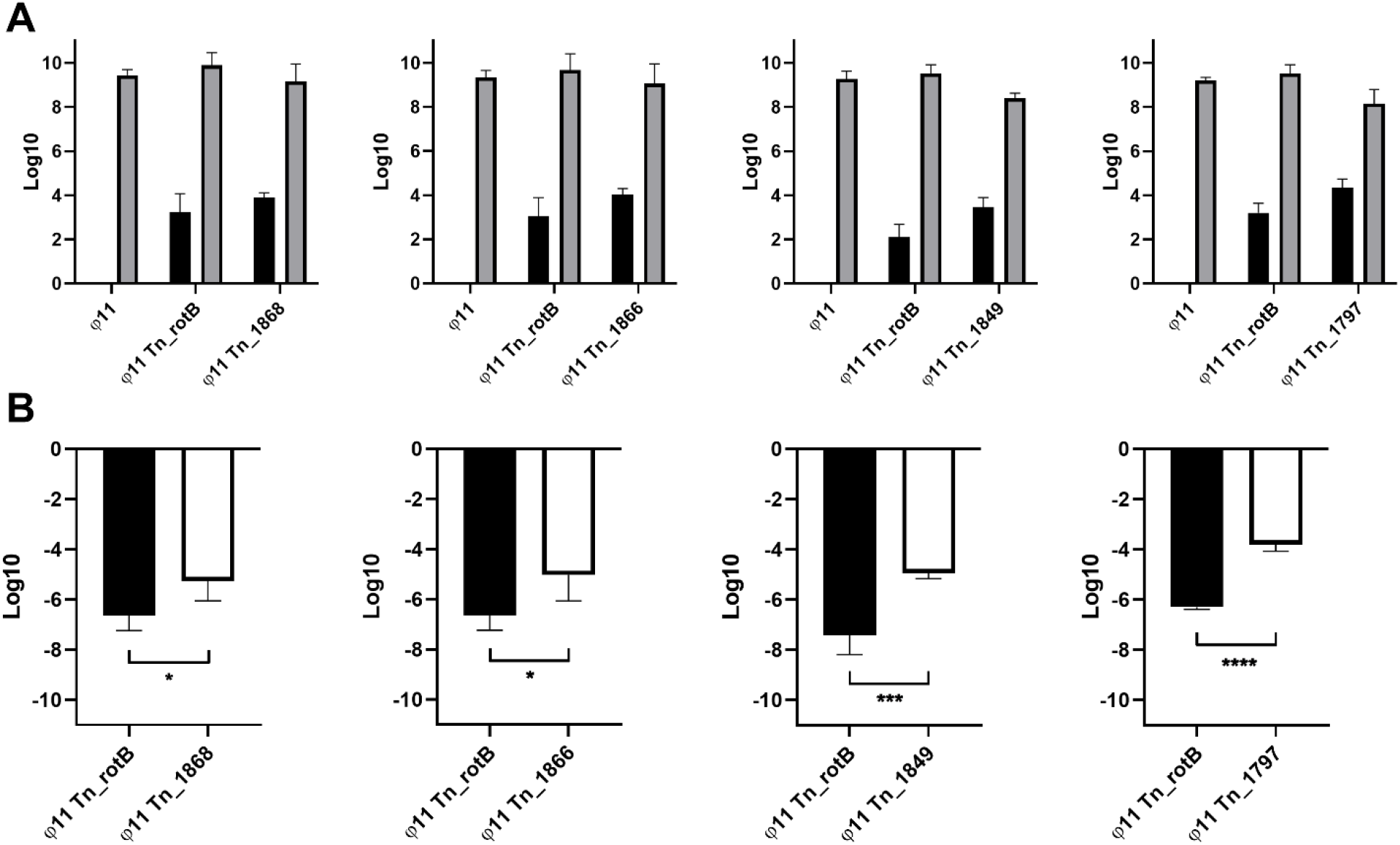
The CFU ml^-1^, PFU ml^-1^, and CFU/PFU transduction frequency of the transposon mutants located up- and downstream of the φ11 integration site. **A** shows the CFU ml^-1^ (black) and PFU ml^-1^ (grey) for control strains 8325-4 φ11 and 8325-4 φ11 Tn_rotB with the 4 different Tn mutants up- and downstream of the φ11 integration site (Tn inserted into SAUSA300_1868, SAUSA300_1866, SAUSA300_1849, and SAUSA300_1797 respectively). The values shown have been log transformed and show the mean and standard deviation (SD) for 4 biological replicates. **B** shows the CFU/PFU transduction frequency for each mutant compared with Tn_rotB (mean and SD, 4 biological replicates). Unpaired t tests were performed and p-values from left to right were 0.0312*, 0.0346*, 0.0008***, <0.0001****.

### Transposon mutants downstream of the φ11 integration site are transferred by lateral transduction

Next was to further define the type of transduction occurring with these 4 chromosomal transposon mutants. Specialized transduction occurs in the immediate flanks neighbouring the prophage, so it was not a likely explanation for Tn_1797. Lateral transduction occurs in bacterial DNA up to 100 kb downstream of a laterally transducing prophage, based on the pac site encoded by that phage. In the case of φ11, the pac site is located in the small terminase gene and this means the region downstream of the att_R_ (<1.923 Mb) is transferred by lateral transfer, whilst the region upstream of the att_L_ (>1.967 Mb) is not (14). To establish whether the higher transduction frequencies seen in the mutants with transposons located downstream of φ11 were due to the effects of the specific transposon insertions or the more general effect of lateral transduction, we tested the transduction frequency of other transposon mutants from the same region of DNA. We used three different mutants where the transposon was situated close to the downstream transposon insertion sites identified from our pooled screens, but had not been identified by the screen (Tn_1799, Tn_1843, and Tn_1852). We transduced these transposon mutants into the 8325-4 φ11 background and measured the transduction frequency following induction with mitomycin C. All three of the transposon mutants had a significantly higher transduction frequency than the Tn_rotB control (Fig 4), and were comparable to the Tn_1849 and Tn_1797 mutants. That these mutants also had a higher transduction frequency similar to the mutants from the screen, suggested that the cause for these higher rates was lateral transduction. Since lateral transduction only occurs when a prophage is induced, we tested the transduction frequency of both the original mutants and these ‘neighbouring’ mutants in an infection set up, where lateral transduction could not occur.

**Figure 4.**
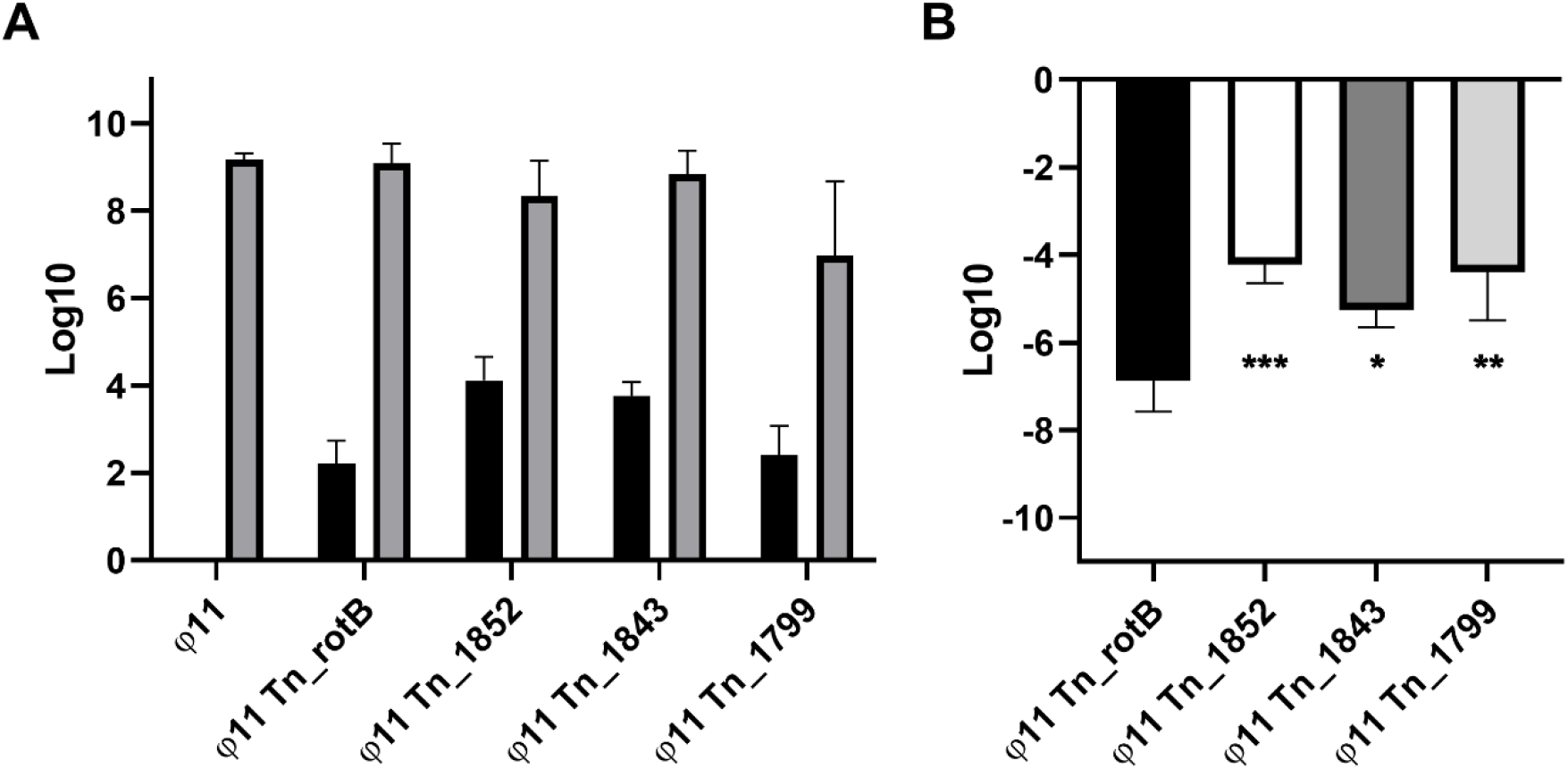
The CFU ml^-1^, PFU ml^-1^, and CFU/PFU transduction frequency of transposon mutants from the chromosomal region downstream of the φ11 integration site that were not selected in the original transduction screen. **A** shows the CFU ml^-1^ (black) and PFU ml^-1^ (grey) for the controls 8325-4 φ11 and 8325-4 φ11 Tn_rotB with 3 different Tn mutants inserted separately in genes downstream of the φ11 integration site (Tn inserted into SAUSA300_1852, SAUSA300_1843, and SAUSA300_1799 respectively). The values shown have been log transformed and show the means and standard deviation for 4 biological replicates. **B** shows the CFU/PFU transduction frequency (mean and SD, 4 replicates) for each mutant compared with Tn_rotB. A one-way ANOVA was performed with Dunnetts multiple comparison test comparing the mean of the Tn_rotB control with each mutants mean, and p-values from left to right were 0.0006***, 0.0203*, and 0.0010**.

To measure the transduction frequency following infection rather than induction, the transposon mutants in the original JE2 library background were infected with φ11 and the transduction frequency of the resulting lysate was established. The transduction frequencies of all the transposon elements located downstream of φ11were higher than the Tn_rotB control (Fig 5A-D); however, these transduction frequencies were reduced when compared to the previous induction-linked frequencies (Fig 5E-G). This supported the idea that lateral transduction was the main cause of the increased transduction frequencies downstream of the φ11 integration site.

**Figure 5.**
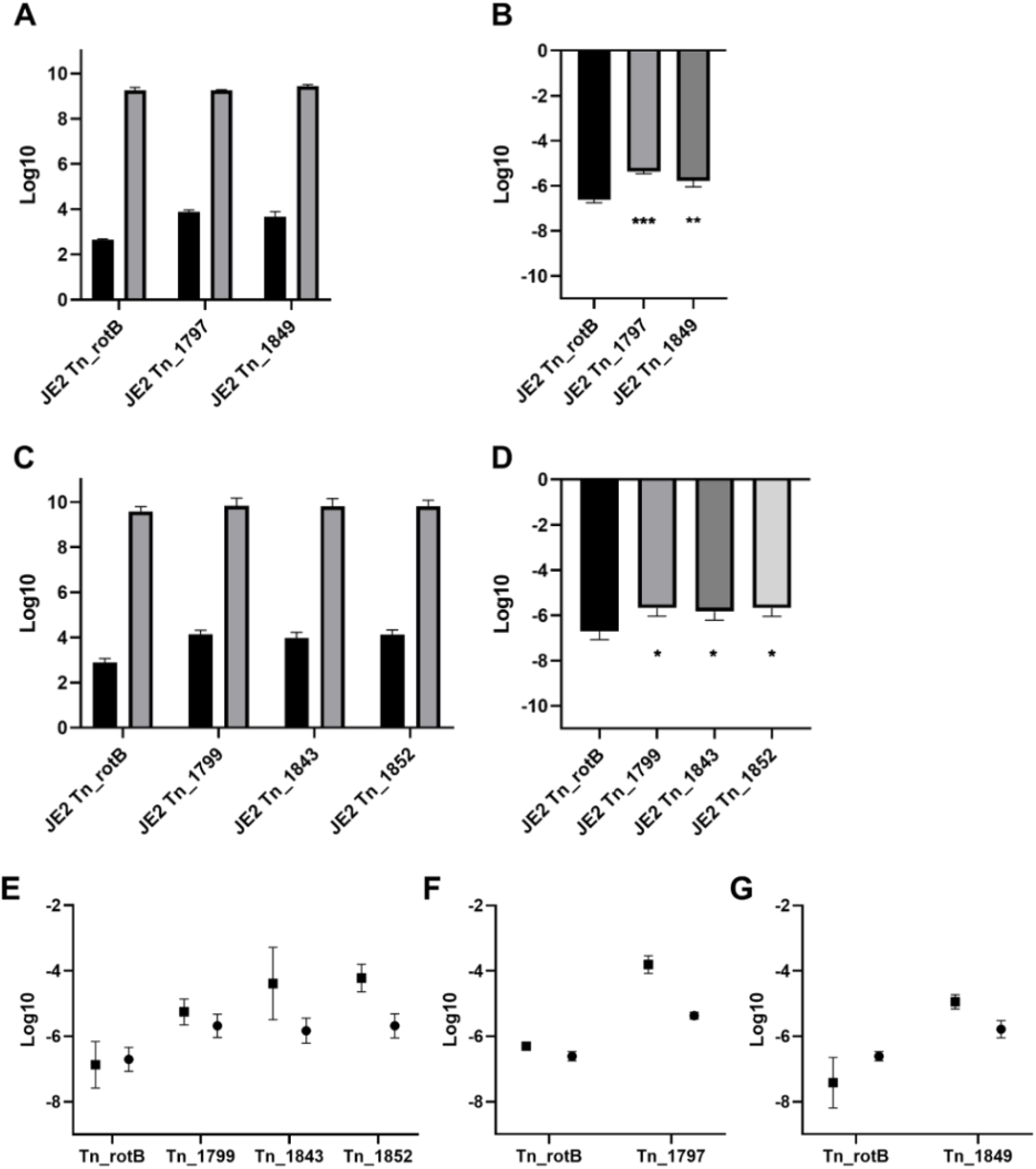
The CFU ml^-1^, PFU ml^-1^, and CFU/PFU transduction frequency of downstream JE2 transposon mutants following infection with φ11. **A** shows the CFU ml^-1^ and PFU ml^-1^ for the control JE2 Tn_rotB and the different screen identified Tn mutants downstream of the φ11 integration site following infection with φ11 (Tn inserted into SAUSA300_1797 and SAUSA300_1849 respectively). **B** shows the CFU/PFU transduction frequency for each mutant compared with Tn_rotB. P-values for a one-way ANOVA with Dunnetts multiple comparison test were 0.0003*** and 0.0027** (left to right). **C** shows the CFU ml^-1^ and PFU ml^-1^ for the control JE2 Tn_rotB and the other Tn mutants downstream of the φ11 integration site following infection with φ11 (Tn inserted into SAUSA300_1799, SAUSA300_1843 and SAUSA300_1852 respectively. **D** shows the CFU/PFU transduction frequency for each mutant compared with Tn_rotB. P-values for a one-way ANOVA with Dunnetts multiple comparison test were 0.0233*, 0.0476*, and 0.0233* (left to right). All values shown have been log transformed and show the means and SD from the 3 replicate values. **E-G, the CFU/PFU transduction frequency following either infection or induction with φ11. E** compares the induction (square) and infection (circle) transduction frequencies of the neighbouring transposon mutants and the Tn_rotB control, showing the means and SD. P-values for a two-way ANOVA with Sidak multiple comparison test were 0.9944^ns^, 0.8400^ns^, 0.0212* and 0.0190* left to right. **F** and **G** compare the induction (square) and infection (circle) transduction frequencies of the Tn_rotB control together with Tn_1797 and Tn_1849, respectively. P-values for a two-way ANOVA with Sidak multiple comparison test were **(F)** 0.0830^ns^ and <0.0001**** **(G)** 0.0874^ns^ and 0.0747^ns^.

### Transposon mutants upstream of the φ11 integration site are transferred by specialised transduction

To establish whether the genes located upstream and close to the φ11 integration site were transduced at higher frequencies due to gene specific effects or because of specialised transduction, additional transposon mutants from this region were identified and tested. The mutants in the same region as Tn_1866 and Tn_1868 showed comparable transduction frequencies, whilst the mutant furthest away from the phage integration site showed a gradual reduction in transduction frequency, albeit higher than the Tn_rotB control (Fig 6. Tn_1865, Tn_1883 and Tn_1898 vs Tn_1916, respectively).

**Figure 6.**
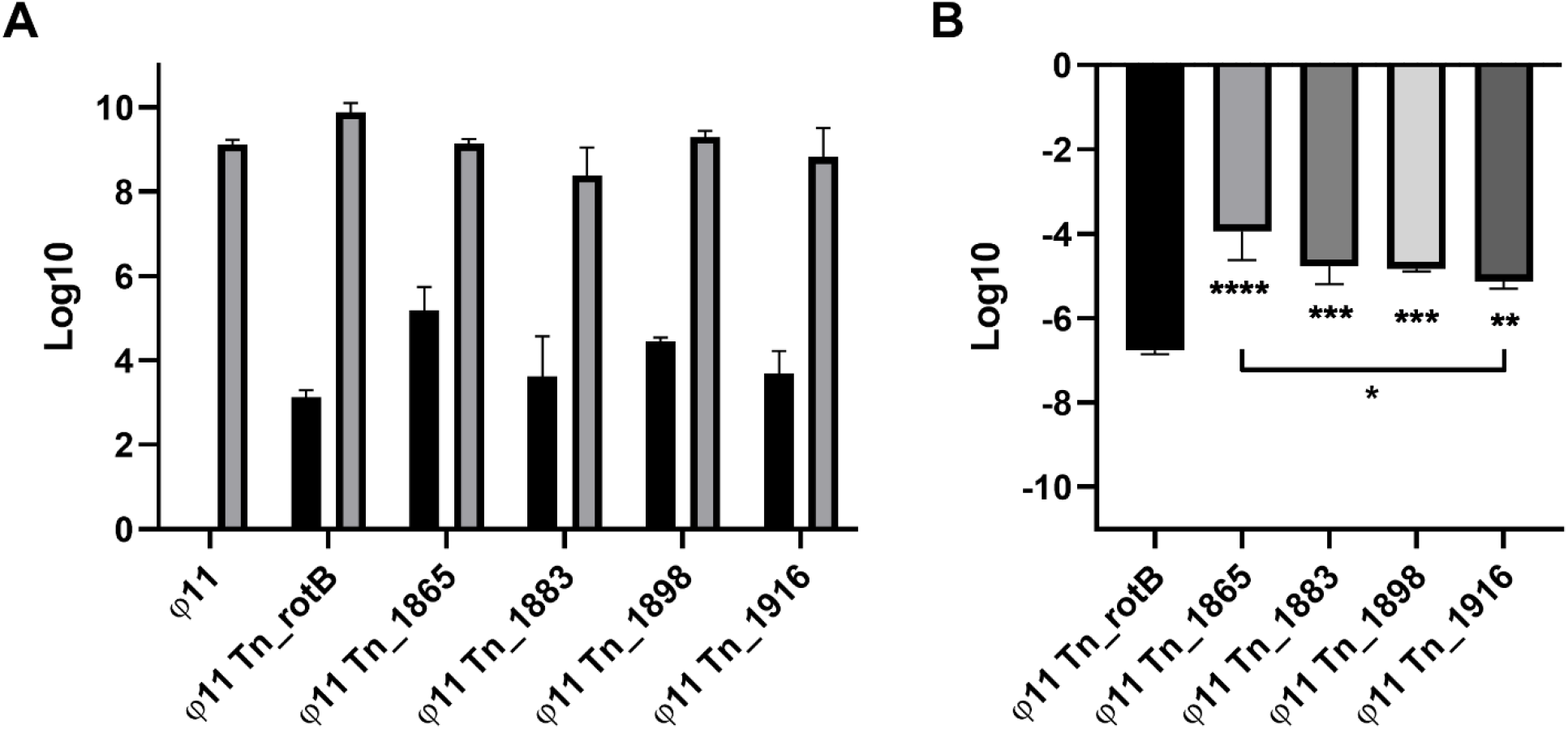
The CFU ml^-1^, PFU ml^-1^, and CFU/PFU transduction frequency of transposon mutants from the chromosomal region upstream of the φ11 integration site that were not selected in the original transduction screen. **A**, shows the CFU ml^-1^ and PFU ml^-1^ for the controls 8325-4 φ11 and 8325-4 φ11 Tn_rotB with 4 different Tn mutants inserted separately in genes upstream of the φ11 integration site (Tn inserted into SAUSA300_1865, SAUSA300_1883, SAUSA300_1898 and SAUSA300_1916 respectively, progressively getting further away from the φ11 insertion site from left to right). **B**, shows the CFU/PFU transduction frequency for each mutant compared with Tn_rotB. A one-way ANOVA was performed with Tukey multiple comparison test comparing the means. Comparison of the Tn_rotB control with each mutants mean gave p-values from left to right of <0.0001 ***, 0.0005 ***, 0.0006***, and 0.0022*. An additional comparison of the Tn mutant closest to the φ11 insertion site (Tn_1865) to the one furthest away (Tn_1916) is indicated by the black line, with a p-value of 0.0188*. All values shown have been log transformed and show the means and SD for 3 biological replicates.

These results suggested that specialised transduction was occurring in the region immediately upstream of the φ11 and causing the higher levels of transduction seen. Furthermore, when the equivalent JE2 transposon mutants were tested following φ11 infection, the transduction frequencies were lower and not significantly different to the Tn_rotB control (Fig 7). Given that specialised transduction requires integration of the phage genome into the host chromosome, which isn’t thought to occur during infection, this further supports that the upstream transposon mutants were being identified due to the occurrence of specialised transduction. Indeed, by using primers annealing within the NTML transposon (Martn-ermR) and the φ11 phage (phi11-direction-3f) it was possible to amplify DNA products from induced and precipitated lysates of 8325-4 φ11 Tn_1866 and 8325-4 φ11 Tn_1868 (Fig. 8). This indicated that some particles contained the upstream regions of bacterial DNA alongside the N-terminal of the phage genome, as would be expected in cases of specialised transduction.

**Figure 7.**
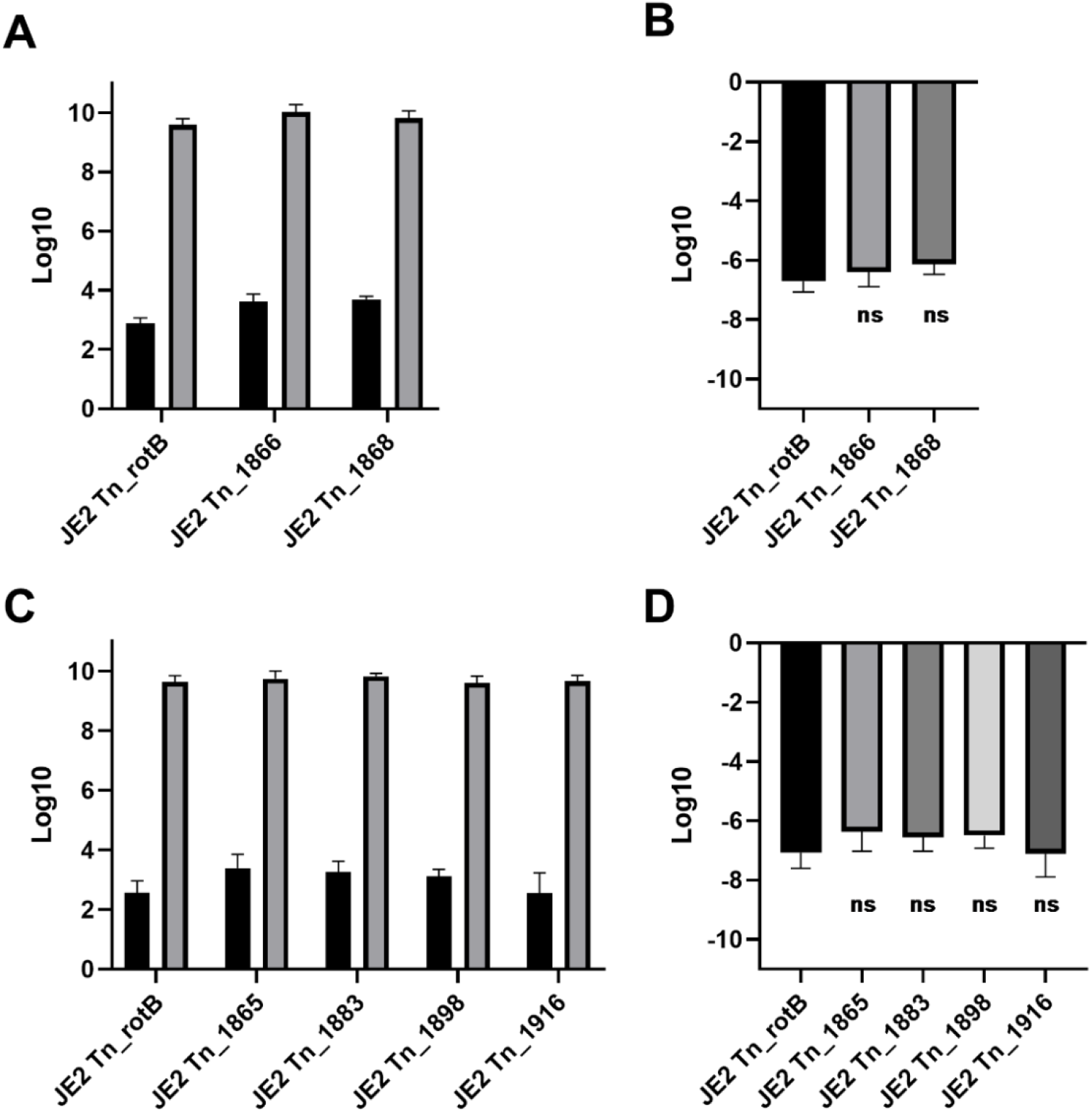
The CFU ml^-1^, PFU ml^-1^, and CFU/PFU transduction frequency of upstream JE2 transposon mutants following infection with φ11. **A**, shows the CFU ml^-1^ and PFU ml^-1^ for the control JE2 Tn_rotB and the different screen-identified Tn mutants upstream of the φ11 integration site following infection with φ11 (Tn inserted into SAUSA300_1866 and SAUSA300_1868 respectively). **B**, shows the CFU/PFU transduction frequency for these Tn mutants compared with Tn_rotB. P-values for a one-way ANOVA with Dunnetts multiple comparison test were 0.5807^ns^ and 0.2225^ns^. **C**, shows the CFU ml^-1^ and PFU ml^-1^ for the control JE2 Tn_rotB and the different neighbouring Tn mutants upstream of the φ11 integration site following infection with φ11 (Tn inserted into SAUSA300_1865, SAUSA300_1883, SAUSA300_1898 and SAUSA300_1916 respectively). **D**, shows the CFU/PFU transduction frequency for the ‘neighbour’ Tn mutants compared with Tn_rotB. P-values for a one-way ANOVA with Dunnetts multiple comparison test were 0.4197^ns^, 0.6682^ns^, 0.5882^ns^ and 0.9999^ns^ (left to right). All values shown have been log transformed and show the means and standard deviation from the 3 replicate values.

**Figure 8.**
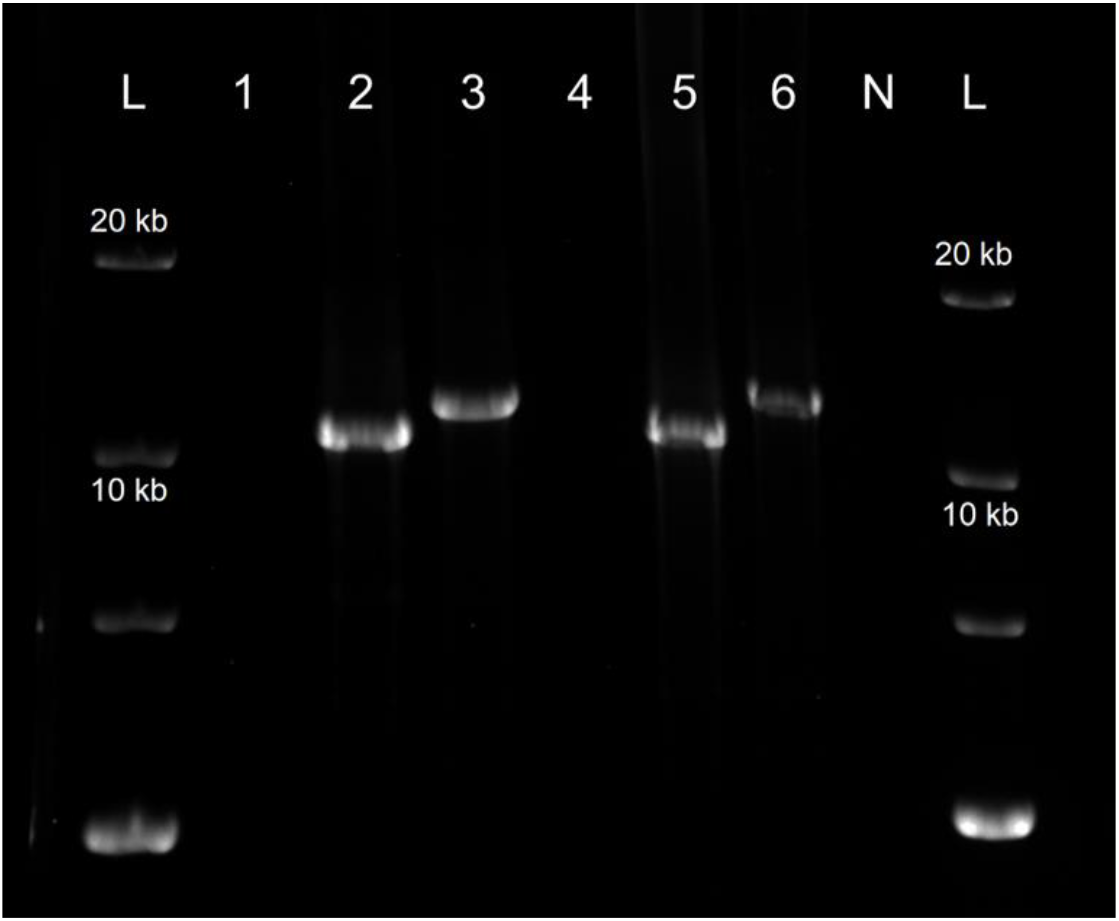
Amplified specialised transduction fragments from precipitated phage DNA. A 0.6% agarose gel showing the PCR product of a long-range amplification using Takara LA polymerase. Templates were as follows, 1. DNA extracted from precipitated phage capsids from an induced lysate of 8325-4 φ11 Tn_RotB, 2. DNA extracted from precipitated phage capsids from an induced lysate of 8325-4 φ11 Tn_1866, 3. DNA extracted from precipitated phage capsids from an induced lysate of 8325-4 φ11 Tn_1868, 4. Bacterial DNA from a culture of 8325-4 φ11 Tn_rotB, 5. Bacterial DNA from a culture of 8325-4 φ11 Tn_1866, 6. Bacterial DNA from a culture of 8325-4 φ11 Tn_1868, N. No template. The expected size for region φ11 to Tn_1866 was ~10.6 kb and ~ 11.7 kb for region φ11 to Tn_1868. Three biological replicates were performed including induction, precipitation, DNA extraction and PCR, and the above gel is representative.

### Regions containing the preferentially transduced transposon mutants are packaged as transductants at higher rates than the rest of the genome

In addition to utilising our NTML transduction screen to identify transposon mutants that were preferentially transduced, in a separate experiment we induced the φ11 in strain 8325-4 φ11 with mitomycin, and sequenced the resultant, precipitated phage φ11 lysates to identify any bacterial DNA that had been packaged in phage capsids. Fig. 9 shows the mapping of the reads to 8325-4, thus removing the majority of reads mapping to the phage and allowing the lower frequency reads that mapped to the bacterial genome to be seen. The regions up- and downstream of the φ11 attB (indicated by the red line) had higher numbers of mapped reads than the other regions of the genome, supporting our screening data in which the identified mutants were clustered in this same region. The pattern of the downstream bacterial DNA indicates that lateral transduction may be responsible, with a gradual reduction of mapped reads from high numbers near to the attB, decreasing the further away from the attachment site. Lateral transduction has been shown to taper off in this fashion, whilst still facilitating transfer of ~ 200 kb of downstream bacterial DNA (14). Here we see slightly elevated read mapping as far away as 798 kb from the attB site. The read alignments of the upstream bacterial DNA encompass 117 kb of DNA upstream of the attB. These reads could represent the effect of specialised transduction following the incorrect excision of the phage, albeit representing a larger region than one would expect with a phage genome length of ~45 kb.

**Figure 9.**
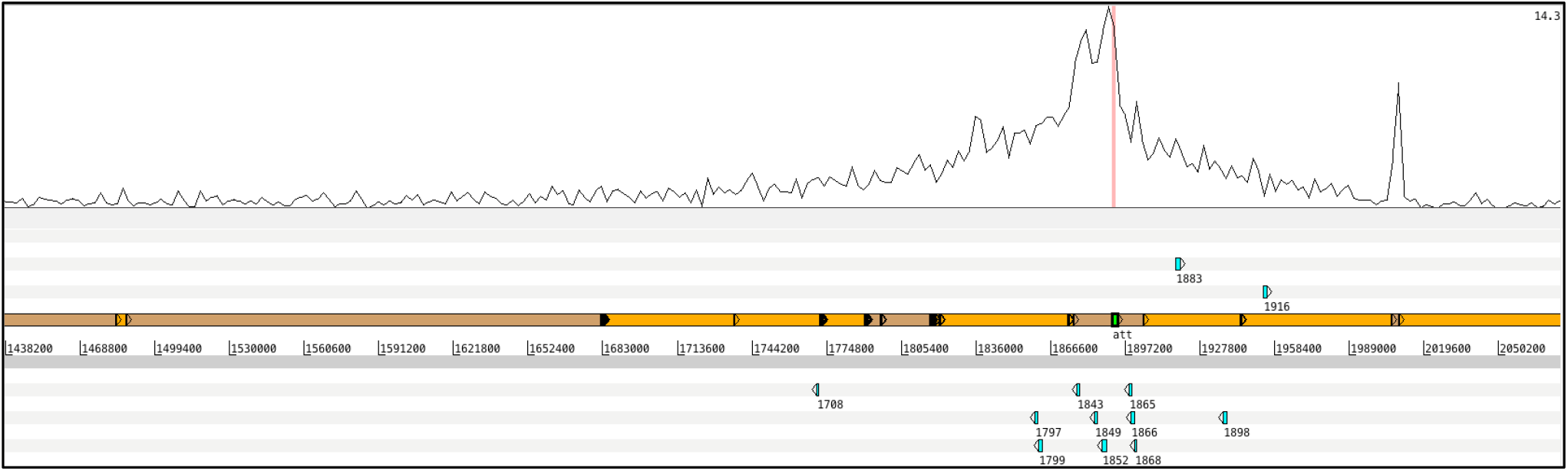
Mapping of lysate-derived DNA to the 8325-4 chromosome. The top graph shows read mapping to part of the 8325-4 chromosome between 1.44 Mb and 2.05 Mb. The red line indicates the site of insertion of φ11 in the lysogen. Beneath this is the six frame translation (top three and bottom three lines) where the genes referred to in this study are represented by blue boxes, with the NTML gene numbers included. The central two lines represent the forward and reverse DNA strands, with the concatenated assembly contigs identified with alternating orange and brown boxes. The narrow peak of read mapping to the right of the insertion site is caused by multiple mapping to a ribosomal RNA operon.

## Discussion

This paper describes a high throughput method by which the mobile regions of bacterial genomes can be identified. By screening pooled transposon mutants, we were able to identify different modes of phage-related horizontal gene transfer, including phage transfer, lateral and specialised transduction. The method gave a manageable number of transposon mutants to further investigate and could be applied to any strain for which there is a transposon mutant library available. Prior to our screen we had anticipated obtaining two types of mutants, namely those where the transposon was placed in the genome such that it could be transferred with higher frequency, for example by lateral transduction, and mutants that due to inactivation of a bacterial gene would lead to increased transduction efficacy. The results showed, however, that we primarily obtained mutants belonging to the first category and, in addition, transposon elements that had inserted into mobile genetic elements. Thus, for four of the resulting transposon mutants, the transposon element was inserted in phages being endogenous to the JE2 strain, namely the USA300φSa2 and USA300φSa3, and these had been transferred to the 8325-4 background. This phage transfer is not totally unexpected, since prophages can be induced by an infecting phage (34) and the 8325-4 strain encodes attachment sites for these phages. However, it is interesting to see both phages being successfully transferred and the resulting lysogens predominate their pools. It suggests that the frequencies of prophage induction, during infection or by spontaneous release, combined with lysogenization is greater than the transduction frequencies in this setup. One surprising transfer event is that of the transposon inserted in gene SAUSA300_0810 from JE2 SaPI5, which is not present in the 8325-4 φ11 recipient, but the transposon was able to transfer to this strain anyway. On closer inspection, it appears that this could be a case of generalised transduction, as there was no evidence of the SaPI5 integrase that would indicate whole island transfer (data not shown) and NCBI Blast comparisons identified a region close to the transposon insertion with homology to a chromosomal region in 8325-4. Further investigations would have to be done to confirm exactly how and where this transduction took place, but it shows that transfer can occur in a multitude of ways and that our methodology captures these.

For two of the transposon mutants that were preferentially selected in our pooled screens, the transposon elements were located downstream of the φ11 attachment site (SAUSA300_1849 and SAUSA300_1797). These mutants displayed high transduction frequencies when transferred following induction of a φ11 lysogen, compared to a control where the antibiotic resistance marker was located elsewhere in the bacterial genome, and this effect was reduced when transfer was monitored following infection of the strains by φ11. These results indicate that the increased transduction was dependent on induction of an integrated φ11, which is one of the prerequisites for lateral transduction (15). Although both specialised and lateral transduction require integration, specialised transduction is limited to regions immediately neighbouring the integrated prophage. Therefore, though both specialised and lateral transduction could potentially transfer Tn_1849, which sits close to the prophage (~8000 kb), Tn_1797 sits >39 kb downstream of the φ11 prophage, nearly a full phage genome length away, so is unlikely to be transferred by specialised transduction. Furthermore, the original study reporting lateral transduction found that a marker 5 kb downstream of the integrated phage was transferred solely by lateral transduction and the researchers did not see any specialised particles containing this region, suggesting all our downstream transfer can be attributed to lateral transduction (15, 17). This notion is supported by the pattern of bacterial DNA sequences obtained from the sequencing of induced phage lysates that indicates extensive packaging of bacterial DNA downstream of the integrated phage. Interestingly, this data actually implicates lateral transduction at an even greater distance downstream of the attB site then has previously been suggested (14), with slightly elevated reads mapping up to 798 kb downstream.

It has been shown that temperate phages are capable of both generalised and specialised transduction (14, 17, 20) but until now, specialised transduction had not been reported in *S. aureus*. However, for the transposon mutants Tn_1866 and Tn_1868 identified in our screen, the transposon element was located immediately upstream of the φ11 integration site at a distance of 7.6 and 8.7 kb, respectively, thus placing it comfortably within a region where specialised transduction could be expected to occur. Further, the increased transduction frequencies of Tn_1866 and Tn_1868 were eliminated when lysates were created by infection rather than induction. Equally, they should not be transferred by lateral transduction, which has been shown to only occur downstream of the phage pac site, in a unidirectional manner defined by the phage packaging machinery (14). This left specialised transduction as the remaining, feasible explanation. Indeed, we were able to identify PCR product containing both φ11 and the transposon using precipitated phage lysate from the strains with Tn_1866 and Tn_1868. This indicated that some phage capsids contained both the att_L_ region of the phage together with the upstream flanking region up to 8.7 kb away. Taken together with the reduced transduction frequency following infection rather than induction, this supports that the upstream genes are transferring through specialised transduction.

In the transfer of the transposon elements located upstream of the φ11 attachment site, we observed ~22-fold higher transduction frequencies compared to our control, which would suggest that, although aberrant excision is generally accepted as a rare event, such particles have a high rate of successful transfer. This could be due to their exploitation of the phage integrase for recombination into the recipient host DNA. On the other hand, it could also be that aberrant excision happens at a higher frequency than expected for φ11. Potentially, the *in situ* replication that has been shown to occur both upstream and downstream of the prophage when φ11 has been induced, but has not yet excised, could be contributing to the rate of specialised transduction(17). However, further work is needed to fully establish the specialised packaging mechanism. In either case, the occurrence of specialised transduction is supported by DNA sequencing of the phage particles, which shows elevated levels of this bacterial upstream region up to 117 kb away from the phage in the precipitated phage lysate. Given that specialised transduction is thought to rely at least in part on the phage integrase for recombination, this could contribute to why we only see upstream genes transferred in this way (17). If prophage φ11 went through an aberrant excision event involving the downstream bacterial flank near the att_R_, it would likely lose the integrase in the process, which could make successful recombination into the recipient less likely. However, given that in our set up the wild-type prophage is present in the recipient, there is likely additional factors contributing to the lack of downstream specialised transduction. Kwoh and Kemper discussed the possible variations of specialised transduction in *Salmonella typhimurium* at length, but there could be differences in specialised transduction between this organism and *S. aureus* (17).

Another interesting aspect of our data is the slightly higher transduction frequencies observed upon infection seen for the Tn mutations located downstream of the φ11, which we predict are transferred by lateral transduction. Such an effect could suggest the integration of a small number of phages during the infection cycle, as proposed in Salmonella phage P22 (35). A ‘lyso-lysis’ effect has been shown in *E. coli* where cells infected by multiple phages at once displayed simultaneous phage lysogenic integration into the genome and entry into the lytic cycle (15). Potentially this is occurring during the φ11 infection cycle and leading to low levels of specialised and lateral transduction being possible. Current investigations of the effect of integration during the lytic cycle on transduction are ongoing.

A limitation of the screening set up described here is that we are analysing the mobility of genes related both to phage infection and prophage induction. Future screens could be optimized to specifically address individual aspects of DNA transfer. Another consideration is the co-selection for transposon mutants with greater fitness, that arise because of the necessary incubation periods during the screening. This may explain why within our pooled mutants we identified a limited number of transposon mutants from the highly transduced regions, even though others were present in the pool. The transposon mutants identified in the screen may well represent those of higher fitness relative to the other highly transduced transposon mutants from that pool.

In summary, this study acts as a proof of concept for using transposon mutant libraries to investigate which regions of the bacterial genome are highly mobile. We show phage transfer, generalised, specialised, and lateral transduction between staphylococcal strains. While the set up employed here was used to identify highly transduced genes and chromosomal locations, it may in fact also reflect what happens over time in the real world. Release of φ11 from lysogens promotes acquisition of bacterial DNA by auto-transduction, and resulting transductants will as lysogens be able to repeat the process (27). Horizontal gene transfer underpins the evolution and adaptability of *S. aureus* and approaches such as the one adopted here will help us increase our understanding of which regions are transferred and how.

